# Vigilance & Cognitive Flexibility: The Effect of Martial Arts Experience on Task Switching

**DOI:** 10.1101/2023.05.28.542654

**Authors:** Ashleigh Johnstone, Paloma Marí-Beffa

## Abstract

Martial Arts can be considered both a sport and a therapy, combining physical exercise with intense periods of concentration (similar to meditation). Previous research suggests that both exercise and meditation can lead to improvements in executive functions (i.e., cognitive flexibility) in adult participants. Results from our lab have previously suggested that martial artists show improvements in the alerting attentional network – a measure of vigilance. The current research aims to investigate the impact of Martial Arts training on a task-switching protocol to measure both vigilance and cognitive flexibility in typical adults. Here we recruited adult martial artists with at least two years of experience, and control participants with no experience. Participants had to respond to either the shape or the colour of a figure in pure and mixed blocks to provide measures of mixing costs (sustained vigilance) and switching costs (cognitive flexibility). Results demonstrated martial artists did not differ from controls in the pure block, but displayed improved performance in the mixed block, revealing an improvement in mixing costs (vigilance). These benefits in vigilance mirror those previously found in attentional tasks, providing convergent evidence on the impact of Martial Arts training on vigilance.

## Introduction

Cognitive flexibility has often been defined as the ability to assess a situation and appropriately adapt the cognitive strategies and processes according to the new situation or environment (Dajani & Uddin, 2015). Indeed, Martin and Rubin (1995) suggested that throughout day to day life, people have to make choices about their behaviour and how to respond to situations. They go on to further suggest that cognitive flexibility is the state of willingness we need to adapt our behaviours and choose the most effective response. This is an important function in everyday life; it allows us to multi-task, problem solve, and even work multi-lingually (Ionescu, 2012). It also allows us to ‘adapt and overcome’ when situations do not work out as first planned. Without cognitive flexibility we would not be able to move on past the obstacles life presents us with. For example, a person finding it difficult to begin the search for a new job following redundancy, or older people sticking to a set routine without engaging in new activities, or children sticking only with one colour of vegetable without trying others. Small examples perhaps, but those which can have a longer-term impact on a person’s life.

Vigilance is also known as sustained attention, and allows us to effectively detect stimuli and respond appropriately given the circumstances we find ourselves in (Sanabria et al., 2019; Sanchis, Blasco, Luna, & Lupiáñez, 2020). As with cognitive flexibility, vigilance is essential in our daily lives. We need to be able to rely on our vigilance capacity to be able to cross the road safely, look after children, or avoid burning our dinner.

In this paper we use a task-switching paradigm to assess the effects of martial arts training on cognitive flexibility and vigilance. Understanding activities and behaviours that can enhance these functions can provide valuable knowledge in developing effective interventions for those experiencing cognitive decline or dysfunction.

### Task-switching

Whilst there are several different iterations of task-switching paradigms, one of the most common is the alternating runs paradigm (Rogers & Monsell, 1995). In this paradigm there are two types of block, a pure block and a mixed block. In the pure block participants are asked to complete one task, either task A or task B, for the whole block. For example, a pure block could look like ‘AAAAAA’. This is often followed by a pure block of the other task, such as ‘BBBBBB’. In the mixed block, however, the tasks are often presented as ‘AABBAA’ which produces two types of trial, repeat trials and switch trials. Repeat trials consist of the same task appearing twice consecutively, whereas the switch trials occur when the task switches such as the ‘AB’ portion of the block.

This way of presenting trials in the alternating runs version of the task, adds an element of predictability. This allows the participant to recognise the predictable pattern, and hold it within their working memory to allow preparation for the upcoming trial (Dreisbach, Haider, & Kluwe, 2002; Ruthruff, Remington, & Johnston, 2001). If the participant has an efficient working memory, this can allow for faster and more efficient responses. Marí-Beffa and Kirkham (2014) use a tennis analogy to further explain this. Tennis players typically return to the centre of the court after playing their shot, because the location of the return ball will be unpredictable. This means that regardless of which side the return ball arrives, the player is in the best location to respond – think of this as moving back to respond to a switch cost. However, if the return ball is predictable, and the player knows that the ball will return to the same side of the court, they will stay where they are for the most efficient response. This is analogous to the predictable trials in the alternating runs paradigm – participants can recognise the pattern and hold it within their working memory, to ensure they are making the most efficient responses, rather than always ‘returning to the centre of the court’.

Due to its predictable nature and the participant needing to keep track of where they are in the pattern, the alternating runs paradigm is believed to have greater working memory demands than other task-switching paradigms (Kamijo & Takeda, 2010). Because of this, many researchers include some form of cue which acts as a signal to the participant and reminds them of their place in the pattern. If no cues are present, this means that participants have to be able to keep track of which trials they have previously completed, as well as how the present trial fits into the pattern. This level of vigilance is not sustainable forever, and therefore it is likely that participants will lose track of the pattern at some point, and so the cues guarantee that they can get back on track – although, in this case, we would expect their response to be similar to that of a switch trial even if it is technically a repeat trial.

The task switching paradigm provides two measures, one of mixing costs and one of switch costs (Monsell, 2003). For both of these measures, a larger cost represents poorer cognitive flexibility and concentration.

### Mixing costs

It has previously been suggested that continuous repetition of a task in which the rules often change is more effortful than continuously completing one single task with no changes, as the first situation requires the participant to be able to maintain two sets of rules and processes throughout (Marí-Beffa & Kirkham, 2014). By comparing the outcome of two situations like this, we gain a measure known as ‘mixing costs’ which are the costs associated with mixing different tasks or rules. This means that mixing costs are calculated by comparing outcomes from trials in the pure block to repetition trials in the mixed block.

Traditionally, there have been differences in the way mixing costs have been calculated and assessed, with Kamijo and Takeda (2010) noting that many researchers calculate mixing costs by comparing all pure trials with all of those from the mixed block – therefore combining repeat and switch trials. As they suggest, this means the mixing cost measure can be contaminated by switch trials which are also assessed separately through switch costs. Therefore, calculating the mixing cost by comparing trials from the pure block to repeat trials from the mixed block, avoids this contamination, and provides us with a separate, distinct measure.

Whilst mixing costs have typically been used as a measure of cognitive flexibility alongside switching costs, it has more recently also been described as a measure of vigilance by (Marí-Beffa & Kirkham, 2014). They suggested that when a person is simply repeating a task with no switches, they require a very low level of vigilance to be able to maintain their performance, as responding becomes almost automatic. However, when the possibility of switches arises, the person needs to become more vigilant in order to prepare and appropriately respond.

Much of the research investigating exercise, or Martial Arts specifically, using a task-switching paradigm focuses on switch costs rather than mixing costs. However, if mixing costs do indeed provide a measure of vigilance, we can look at vigilance related research to inform our current hypotheses.

Previous research from our lab has shown that taking part in Martial Arts training is associated with improved vigilance, as evidenced in differences in the alert attentional network between martial artists and non-martial artists (Johnstone & Marí-Beffa, 2018). The alert index is calculated by looking at how people respond endogenously to unexpected events compared to events with exogenous cues and is a known measure of vigilance. It was found that martial artists were better equipped to effectively and efficiently respond to uncued events showing their increased vigilance in comparison to non-martial artists.

### Switch costs

From an experimental cognitive psychology perspective, switch costs from task switching can tell us about a person’s cognitive flexibility. Switch costs are the costs in performance associated with switching between sequential tasks and come from comparing the switch and repeat trials from the mixed block (Marí-Beffa & Kirkham, 2014). This cost comes from the amount of time taken to disengage with the processes needed to deal with the first trial, processing the new trial, and then engaging the new rules and processes needed to respond. As discussed by Marí-Beffa and Kirkham (2014), this is only the case for switch trials, as when the trial is repeated, the participant does not need to change their approach to be able to respond, as they simply need to maintain the rule.

With respect to exercise effects on cognitive flexibility, Masley, Roetzheim, and Gualtieri (2009) recruited participants for an experiment investigating the effects of aerobic exercise on a series of cognitive measures. Participants were asked to complete the CNS Vital Signs neurocognitive battery, which provided measures of memory, attention, information processing speed, and cognitive flexibility. They found an increase in cognitive flexibility which was proportional to the level of aerobic exercise completed by the participants – i.e., moderate or intense. While not a martial arts focused study, this does highlight the potential effects of exercise on cognitive flexibility.

More recently, Lo et al. (2019) investigated the effects of martial arts training on the set-shifting abilities of children, which they explained as the ability to switch between different tasks or goals. Using a task-switching paradigm, they found that long-term judo training (defined as a minimum of four years regular practice) was associated with improved set-shifting compared to control participants with no martial arts experience. Lo et al. (2019) explained this by discussing how the sport trains people to anticipate changes in an uncertain and evolving environment. This training to always try to anticipate the next move and be prepared to change response may enhance the executive functions needed for efficient set-shifting, such as cognitive flexibility and vigilance which leads us to the current experiment.

### Aim and hypotheses

Experiment 1 is a pilot study which aimed to uncover potential confounding demographic variables which could affect the main experimental variables of mixing costs and switch costs. The aim of this was to ensure we could isolate the effects of martial arts experience. Experiment 2 then aimed to investigate the effects of Martial Arts training on vigilance and cognitive flexibility with matched participant groups based on the findings of Experiment 1.

Within Experiment 2, we hypothesised that differences would be most prevalent in mixing costs due to its relationship with vigilance which we have previously found to be influenced by Martial Arts practice (Johnstone & Marí-Beffa, 2018). If this is the case, then it is expected that this would be correlated with years of Martial Arts experience. It is also possible that there will be differences in switch costs between the two groups, due to previous research reporting evidence of changes in cognitive flexibility related to exercise and martial arts.

## Experiment 1

### Method

#### Participants

40 participants were recruited in 2017 from Bangor University’s SONA participant panel system, the demographics of which can be found in Table 1. These were naïve participants that had not taken part in any other study presented in this thesis. They were given an information sheet explaining the purpose of the research, the test settings and the approximate duration. Once they agreed to participate, they were given a consent form containing a summary of the research objectives, approximate duration, full explanation of their right to withdraw at any time without penalty, protocols related to confidentiality, anonymity, password protected storage and sharing of summary group data. Complaints and reimbursement procedures where also included in the consent form. Participants were compensated for their time with course credits. These participants were neurotypical, had normal or corrected to normal vision, and normal hearing. Ethical approval was granted by the Bangor University Ethics and Governance Committee (Ethics Approval #2015-15553).

**Table 1.**
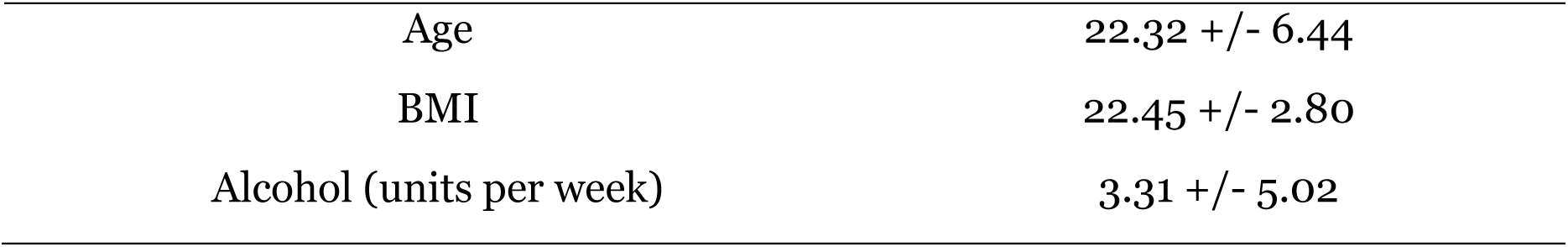
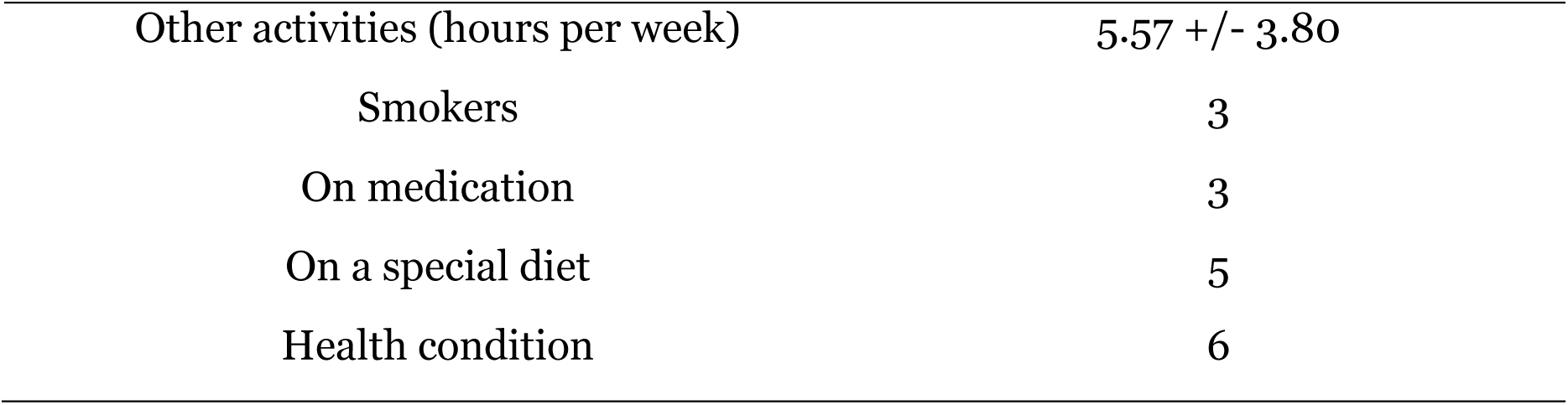
Descriptive data for key participant demographics. Values represent mean averages, +/− standard deviation, or frequencies.

#### Stimuli

Stimuli were presented on screen through EPrime 2.0 [Psychology Software Tools (PST)]. Stimuli consisted of either a square or a circle, presented in blue or red, on a white background. Before these figures appeared, a cue of ‘COLOUR’ or ‘SHAPE’ was presented in black text for 500ms, and then a 250ms blank screen interval. Following this, the target stimulus appeared for 1000ms or until the participant responded, before a 250ms blank screen interval finished the trial.

These target stimuli could appear as either congruent or incongruent, depending on the cue shown. Blue stimuli and square stimuli required a press of the ‘F’ key as a response, whereas red or circular stimuli required a ‘J’ keypress. This means that a blue square (or red circle) would elicit the same response, regardless of whether the participant was asked to respond to the shape or colour of the figure, making it a congruent stimulus. A blue circle (or red square), on the other hand, would be an incongruent stimulus as the required response would depend on the given cue.

#### Design

This study utilised a cued version of the alternating runs task-switching paradigm, which consisted of three blocks (two pure task blocks, and one mixed task block; see Figure 1). All participants completed all three blocks.

**Figure 1.**
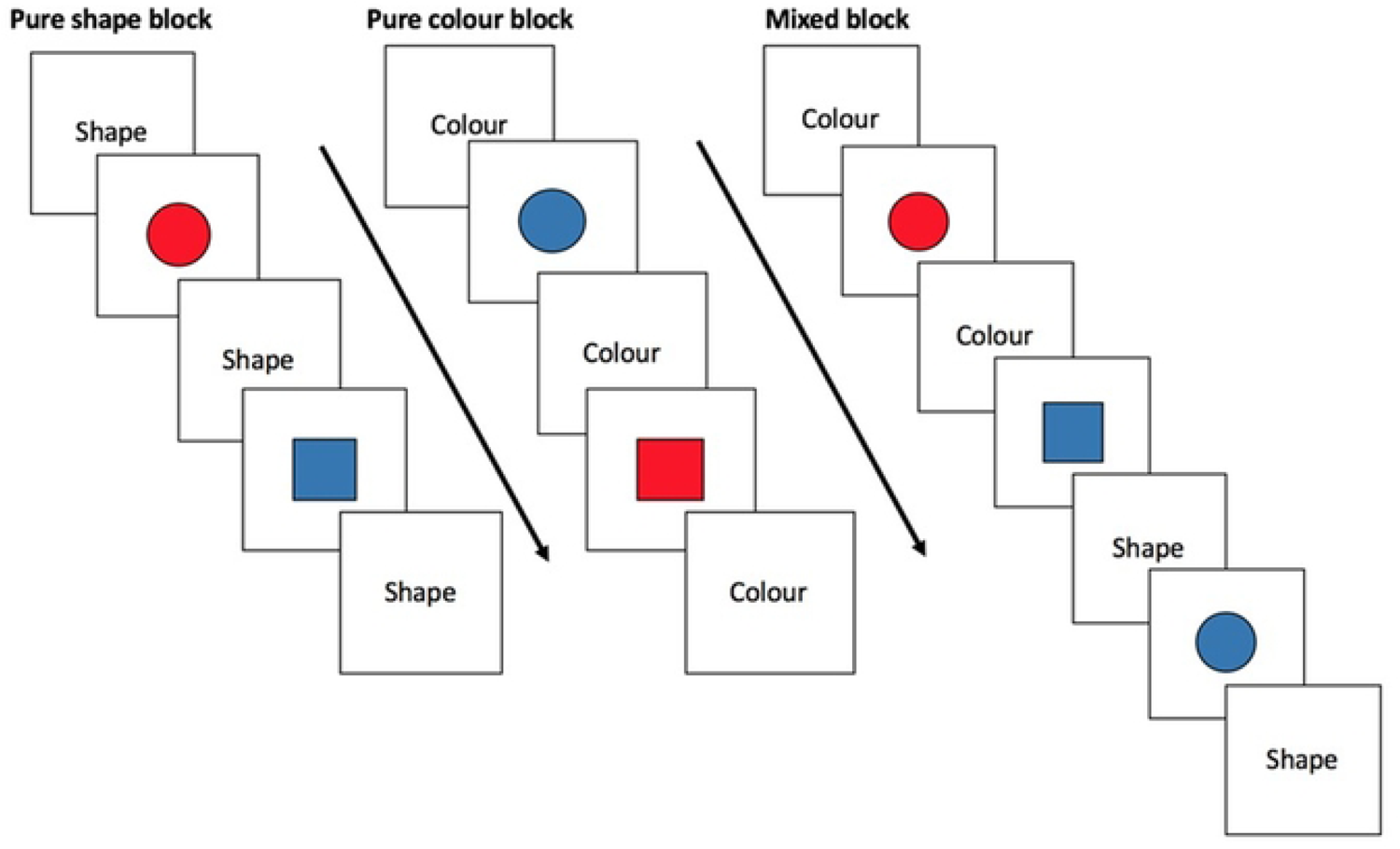
Diagram showing structure of the blocks used in this experiment.

##### Pure Blocks

There were two pure blocks, each consisting of a different task, colour and shape. In one block participants were asked to respond to the colour of the presented figure (i.e., CCCCCC for all trials in the block), and in the other block they responded to the shape of the figure (SSSSSS for all trials in this block). These trials will be referred to as ‘pure trials’ from this point onwards. The order of these blocks was counterbalanced, to ensure that half of the participants completed the colour block first, and half completed the shape block first. This was done to account for any potential contamination effects in the second pure block.

##### Mixed Block

The final block for all participants was the mixed block, which comprised of two alternating trial types. Half of the trials in this block consisted of the participant responding to the colour, or shape for the other half (i.e., CCSSCC). This structure allows us to assess repeat trials and switch trials, (please see Figure 2).

**Figure 2.**
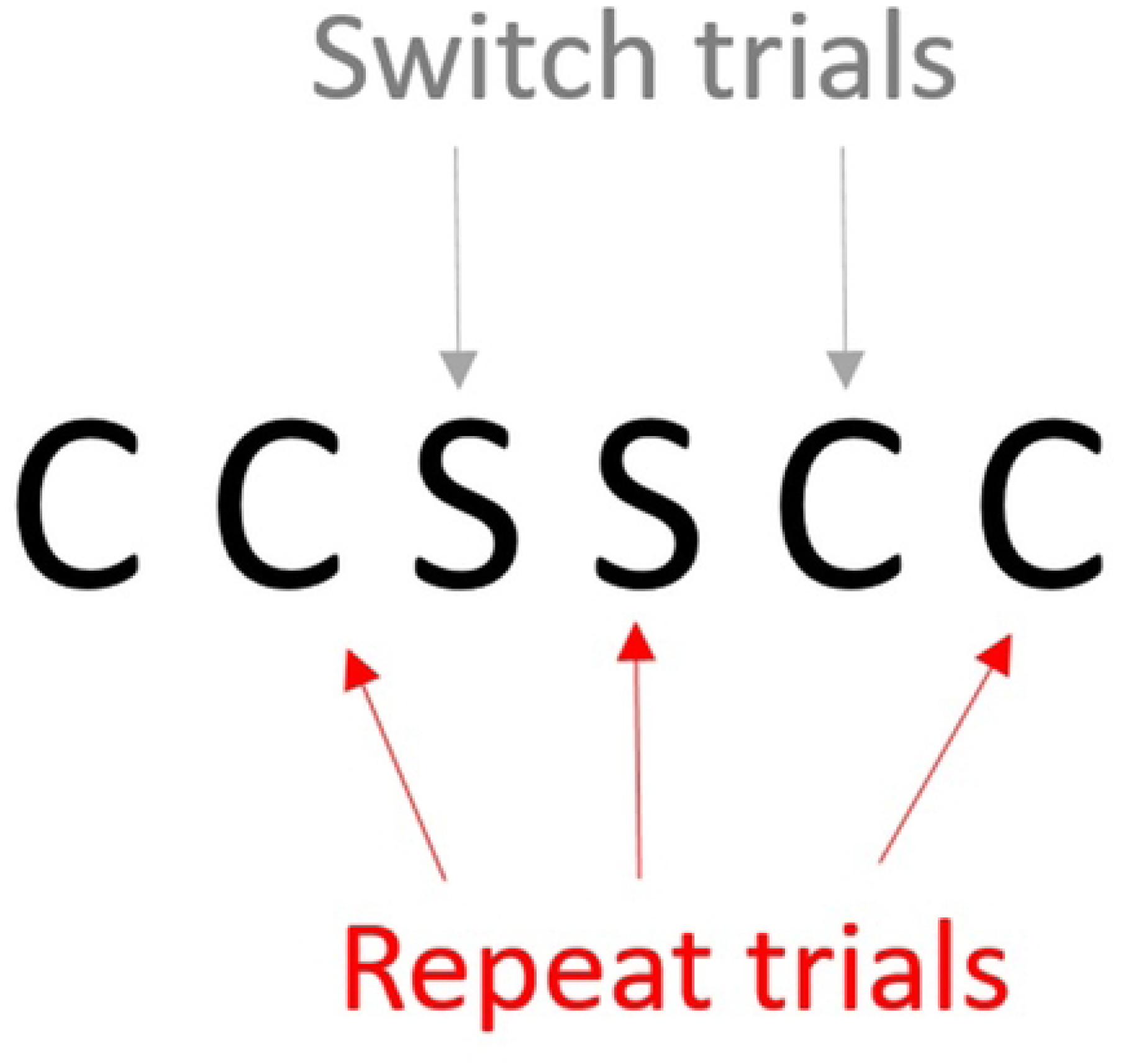
Pictorial representation of order of six trials in the mixed block. ‘C’ represents a colour trial, while ‘S’ represents a shape trial.

#### Procedure

Participants were provided with a consent form, and then asked to complete a demographics questionnaire before partaking in the computer-based task-switching experiment. Before beginning, participants were reminded of the importance of reading the instructions for each block due to the changing instructions throughout the task. They were asked to press the ‘F’ button on a computer keyboard if their response was either ‘blue’ or ‘square’, or the ‘J’ key if their response was ‘red’ or ‘circle’. The two pure blocks contained 144 trials each, and the mixed block consisted of 288 trials (please see Figure 2 for block structure). There were an equal number of congruent and incongruent trials. Participants were not given feedback on their responses throughout the task.

#### Data analysis

Data were pre-processed in EPrime 2.0 to allow for the removal of incorrect trials, and any trials with a RT of under 200ms. This provided an average mean RT for pure, repeat, and switch trials, which were used in the calculations for the mixing costs and switch costs, as described by Kirkham, Breeze, and Mari-Beffa (2012). RT for repeat trials during the pure block were subtracted from RT for the repeat trials from the mixed block to produce the mixing costs. For switch costs, it was calculated as RT for switch trials minus RT for repeat trials from the mixed block. We also ran a series of bivariate Pearson’s correlations to investigate relationships between demographic variables and our experimental variables (mixing and switch costs).

## Results

Significant results from the Pearson’s correlations on the demographic variables and the experimental variables can be found in Table 2 below.

**Table 2.** Pearson correlations between key demographic variables and mixing costs and switch costs.

|  | Mixing Cost | Switch Cost | Age | BMI | Alcohol Units | Other Activities |
| --- | --- | --- | --- | --- | --- | --- |
| Mixing Cost | - | - | - | - | - | - |
| Switch Cost | .161 | - | - | - | - | - |
| Age | -.067 | -.369* | - | - | - | - |
| BMI | -.296 | -.164 | -.032 | - | - | - |
| Alcohol Units | -.004 | -.047 | .355* | .065 | - | - |
| Other Activities | .107 | .163 | -.340* | .102 | -.229 | - |

\*. Correlation is significant at the .05 level (2-tailed)

Age was significantly correlated with switch costs, *r*(40)=-.369, *p*=.019. BMI showed a relationship with mixing costs, although not significantly so, *r*(35)=-.296, *p*=.084. There were also several other correlations between the demographic variables. A significant positive correlation was found between age and units of alcohol per week, *r*(39)=.355, *p*=.026, whilst a significant negative correlation was found between age and hours of reported other activities per week, *r*(40)=-.340, *p*=.032.

## Discussion

From this pilot study we concluded that the age of the participants is significantly associated with their switch costs, and that BMI may be related to mixing costs. We also noted that weekly alcohol units and weekly hours of other activities were correlated with age. This led to the decision to match our martial artist and non-martial artist participant groups on these demographic variables, to avoid confounding effects.

## Experiment 2

Following the pilot study in which key demographic factors were identified, the aim of Experiment 2 was to investigate the effects of Martial Arts practice on mixing costs and switch costs.

### Method

#### Participants

To calculate the optimal sample size, we used G*Power 3.1.9.4 using parameters taken from the pilot study. Using the following stringent criteria, it was calculated that 18 participants in each of the two participant groups would be needed, leading to a total of 36 participants: a correlation of 0.6 between measures, a desired power of 0.95, and an alpha level set to 0.05, to reach an effect size of 0.25 from the required mixed factorial ANOVA.

Ethics procedures were covered by the same ethics approval used in the previous study. A screening questionnaire which assessed Martial Arts experience as well as a range of other demographic factors, was distributed to Bangor University students and non-students from the local community in 2017. Responses to this questionnaire were then used to recruit participants for the experimental phase of the study, leading to a total of 84 participants, 41 in the Martial Arts group, and 43 in the Non-Martial Arts group. Upon recruitment, contact details were destroyed and further data was made anonymous. These participant groups were matched on the key demographic variables identified in Experiment 1. See Table 3 for demographics information for both groups.

**Table 3.** Participant demographics for both participant groups

|  | **Martial Artists** | **Non-Martial Artists** | ***p*** |
| --- | --- | --- | --- |
| *N* (Male) | 41 (18) | 43 (11) | - |
| Age | 23.51 ± 5.82 | 22.19 ± 4.19 | .236 |
| BMI | 23.27 ± 3.35 | 23.86 ± 3.23 | .445 |
| Alcohol (units per week) | 3.70 ± 5.47 | 3.15 ± 4.75 | .627 |
| Smokers | 5 | 3 | - |
| Health Condition | 4 | 3 | - |

Values represent mean averages, ± standard deviation, or frequencies.

The Martial Arts group was made up of participants with experience in the following styles: Multiple (12), Karate (9), Taekwondo (5), Judo (4), Kickboxing (4), Jiu-Jitsu (4), Boxing (2), and Aikido (1).

#### Design and procedure

The task design and procedure were identical to that described for Experiment 1.

#### Data analysis

Again, data were pre-processed in EPrime 2.0 with the same filtering criteria. This provided an average mean RT for pure, repeat, and switch trials, as well as congruent and incongruent trials, for each participant. This meant that each participant had data from six trial types – pure congruent trials, pure incongruent trials, repeat congruent trials, repeat incongruent trials, switch congruent trials, and switch incongruent trials. We calculated global mixing and switch costs (incorporating congruent and incongruent trials) and costs based on congruency. Finally, effect sizes were estimated using partial eta squared, and any correlations were completed using Pearson’s correlations.

## Results

We first analysed all of the data as part of a global ANOVA (2×3×2) including participant group (martial artists and non-martial artist; 2) as between group factor, and switching condition as repeated measures (pure, repeat, switch; 3), and congruency as another repeated measures (congruent, incongruent; 2). Please see Table 4 for descriptive statistics. These main results demonstrated that overall martial artists were 42ms faster than non-martial artists, F(1,82)=4.32, *p*=.041, *η_p_^2^*=.050.

**Table 4.**
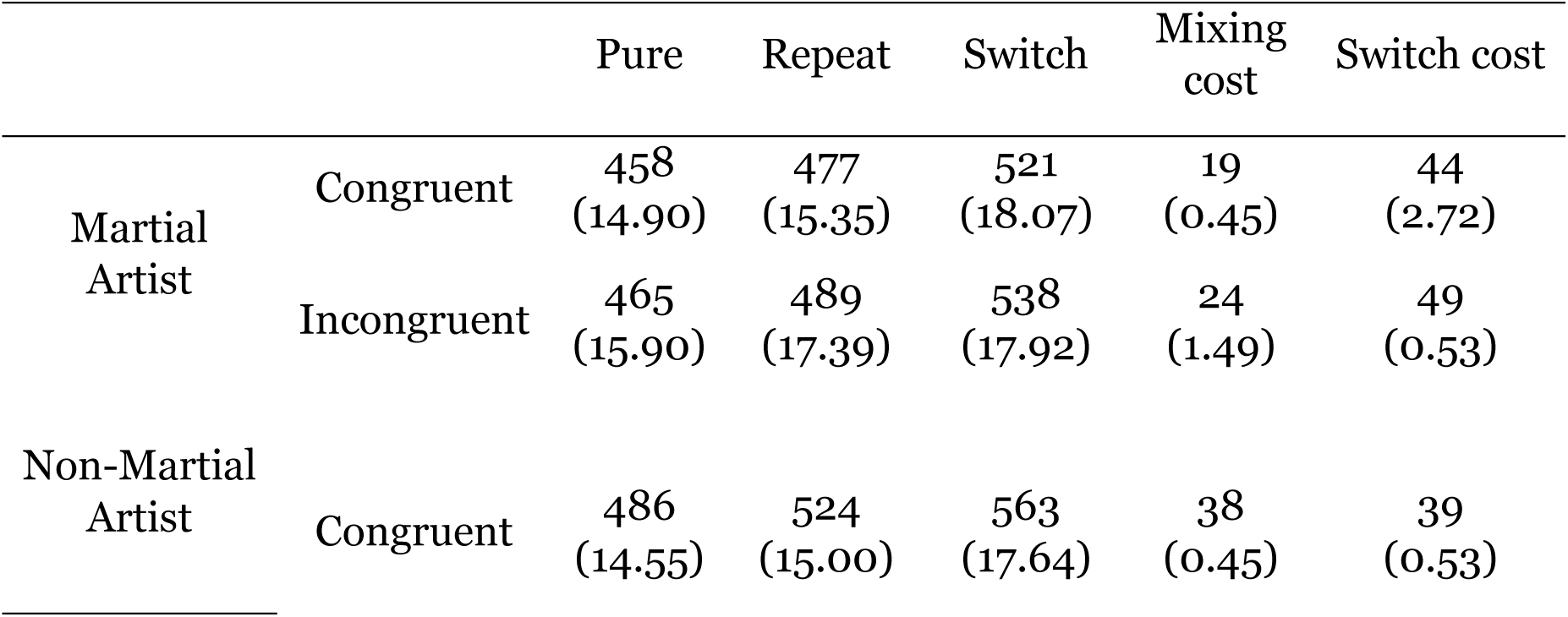
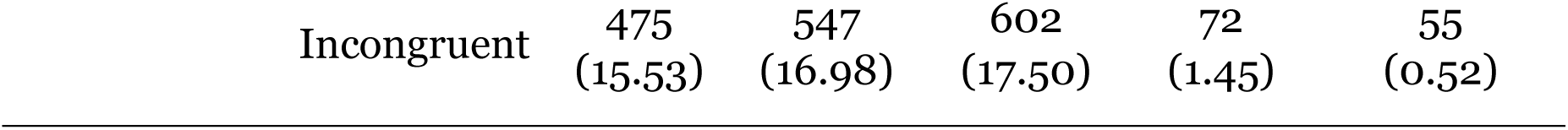
Mean reaction times (in ms) for each trial type, plus mixing cost and switch cost for the two participant groups

There was a main effect of switch condition, F(2,164)=46.15, *p*<.001, *η_p_^2^=*.36, where pure repeat trials were 38ms faster than mixed repeat trials (the mixing cost; t(83)=4.15, *p*<.001). The switch trials were 47ms slower than the repeat trials (switching cost; t(83)=7.72, *p*<.001). There was also a main effect of congruency, F(1,82)=11.95, *p*=.001, *η_p_^2^ =*.127, where congruent trials were 14ms faster than incongruent trials. The switching variable condition changed depending on the congruency, F(2,164)=8.45, *p*<.001, *η_p_^2^ =* .093, with greater differences observed in the incongruent compared to the congruent condition. More importantly, the change of switching conditions as a function of congruency was different across the two groups F(2,164)=4.09, *p*=.018, *η_p_^2^ =*.048.

We followed this interaction by analysing mixing costs and switching costs separately.

### Mixing costs

The data were then submitted to a 2×2×2 mixed ANOVA (Martial Arts group – martial artist, non-martial artist; mixing cost – pure repeat and mixed repeat; congruency – congruent and incongruent). This result confirmed that martial artists were 36ms faster than non-martial artists overall (F(1,82)=3.47, *p*=.066, *η_p_^2^ =*.041), but not significantly so.

In addition, results also confirmed previous findings that the mixing cost was significantly greater (reflecting poorer flexibility) in the incongruent condition than the congruent condition (F(1,82)=7.32, *p*=.008, *η_p_^2^ =*.082). There was a 48ms mixing cost in the incongruent condition that went down to 29ms in the congruent trials. More importantly, this change of mixing cost with congruency was different across martial artist groups (F(1,82)=4.20, *p*=.044, *η_p_^2^ =* .049). The increase of mixing costs for incongruent trials was only observed in the non-martial artist group (F(1,42)=9.93, *p*=.003, *η_p_^2^ =* .191, while no interaction was found with the martial artists (F<1). Non-martial artists demonstrated a 48ms increased mixing cost in incongruent trials compared to martial artists, while they did not differ to martial artists in the congruent trials (see Figure 3).

**Figure 3.**
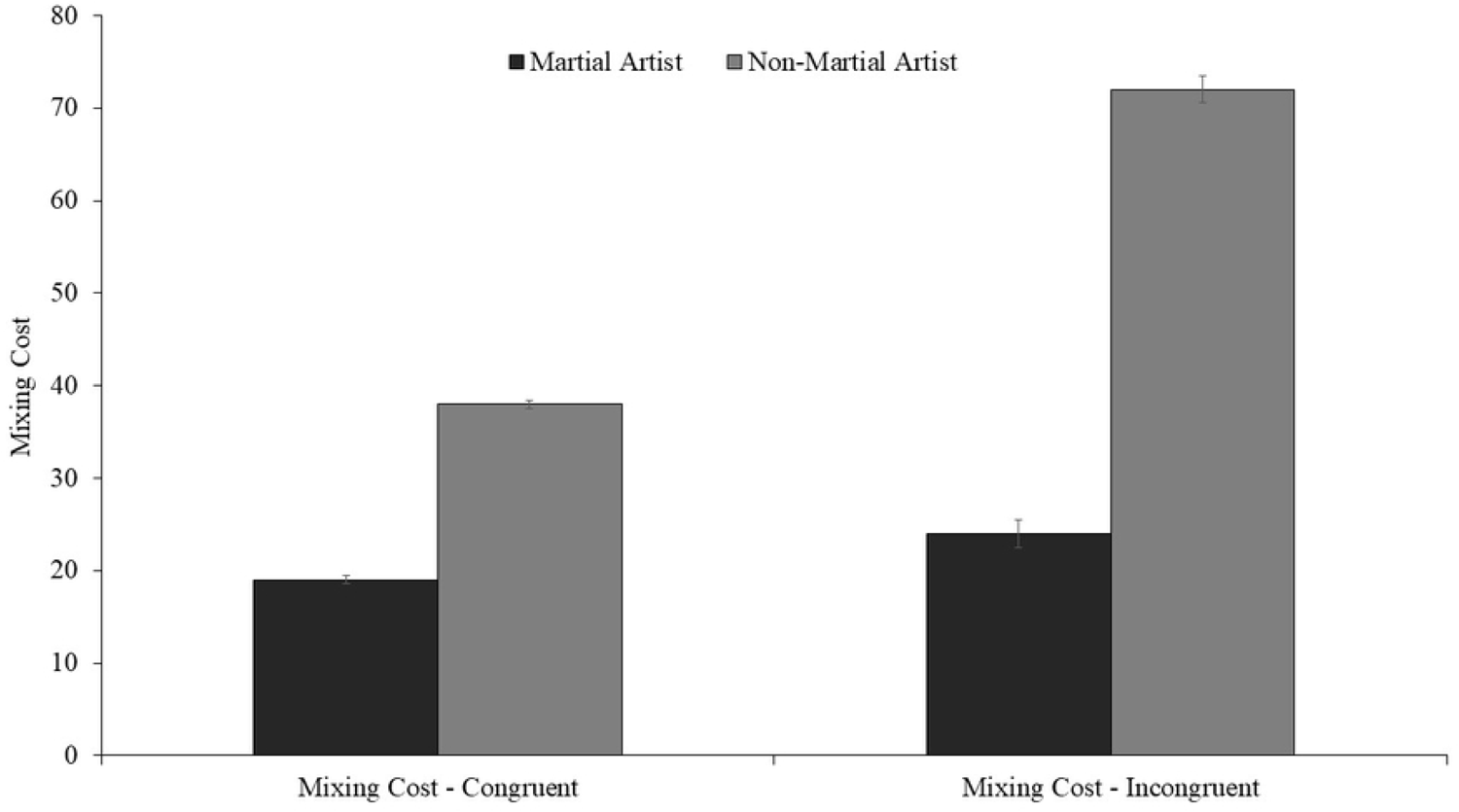
Graph depicting the congruent and incongruent mixing cost, for both participant groups. Error bars represent Standard Error.

Further follow up tests demonstrated no significant differences in the pure repeat conditions, while the only between group differences were observed in the mixed repeat conditions (congruent trials: t(82)=2.2, *p*=.03, incongruent trials: t(82)=2.40, *p*=.019). Importantly, RTs in the incongruent repeat condition significantly correlated with years of MA experience, r(81)=-.245, *p*=.027, but not in the congruent repeat condition, r(81)=-.2, *p*=.074. The correlation between mixing cost in the incongruent trials and MA years was not found to be significant, r(81)=-.21, *p*=.066.

### Switch cost

We analysed the switching costs through a 2×2×2 ANOVA (Martial Arts group – martial artist, non-martial artist; switch cost – switch and mixed repeat; congruency – congruent and incongruent). For this analysis we exclusively considered trials from the mixed block (switch trials and mixed repeat trials). The results demonstrated that martial artists were 53ms faster than non-martial artists overall, F(1,84)=5.48, *p*=.022, *η_p_^2^ =* .063. In addition, congruency effects in the martial artists groups were significantly smaller (14ms) than in the non-Martial Arts group (32ms), F(1,82)=4.45, *p*=.038, *η_p_^2^ =*.051, see Figure 4.

**Figure 4.**
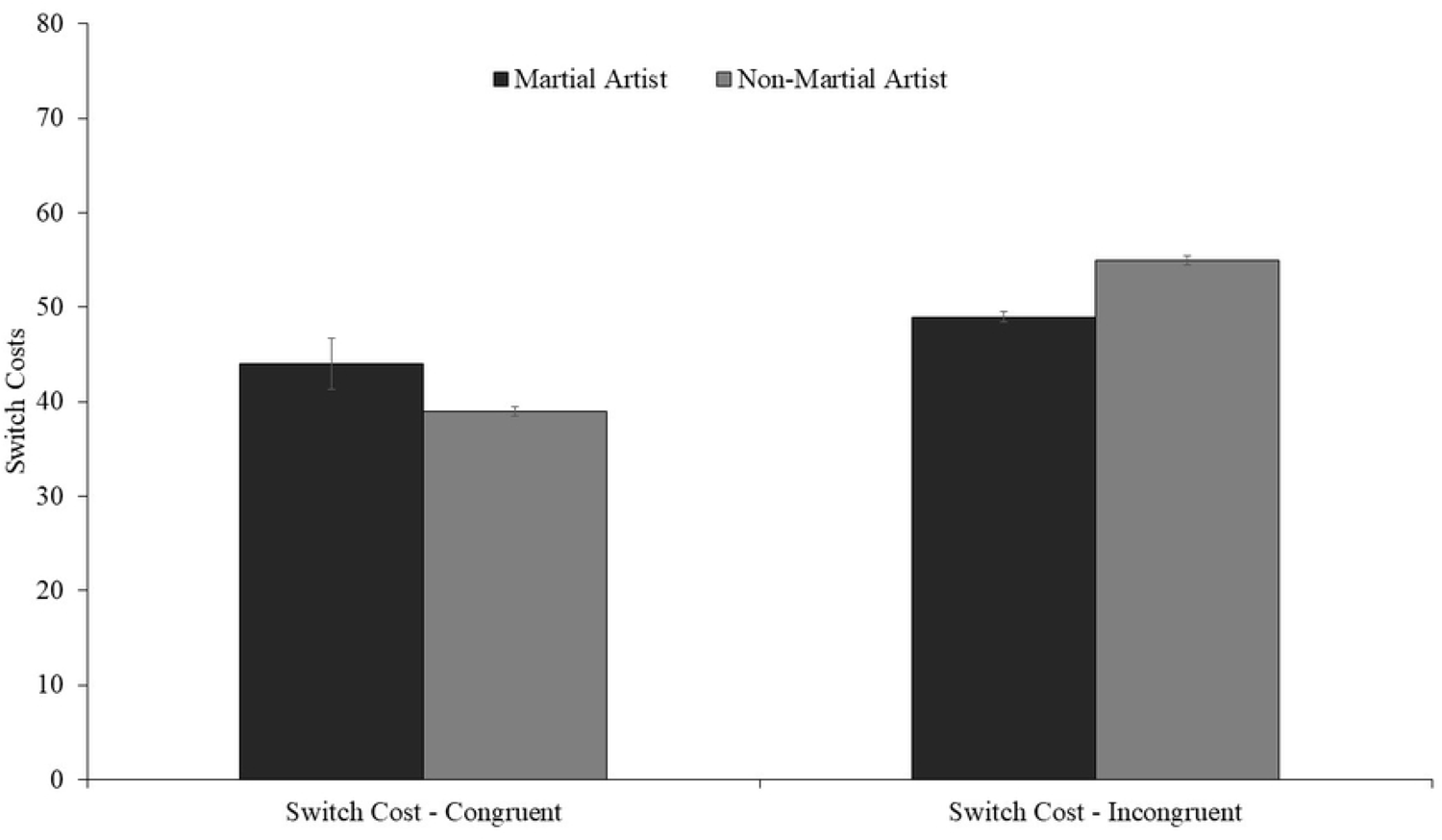
Graph depicting the congruent and incongruent switch cost, for both participant groups. Error bars represent Standard Error.

## Discussion

As predicted, these results suggest a difference in mixing cost associated with Martial Arts practice, with participants from the Martial Arts group performing at a similar level to controls in the pure block, but at a more improved level in the mixed block. Perhaps surprisingly there was no significant difference between the two groups with regards to switch costs.

The finding that martial artists did not differ from controls when considering congruent mixing costs, but performed significantly better than controls for incongruent mixing costs, is interesting. This suggests that the martial artists are less affected by changes in congruency than the non-Martial Arts participants who exhibited a larger mixing cost in the incongruent trials. It is important to remember that a larger cost reflects poorer concentration. We believe this result reflects poorer vigilance from the non-martial artists, and therefore greater vigilance from the martial artists, as the martial artists perform at a similar level regardless of the congruency. This means that the martial artists could still execute an efficient response regardless of whether the stimulus was congruent or incongruent, whereas the non-martial artists were less efficient with incongruent stimuli.

Another possible interpretation of this result is related to working memory demands. The incongruent mixing costs are the element of our paradigm that has the heaviest working memory demand – first, participants have to remember where they are in the pattern of trials to help them predict which trial type will appear next, second, they need to hold the cue in mind, and finally they have to remember the response rules for the different stimuli types to ensure they provide an accurate response. This leads to a heavier working memory demand than any other condition. As discussed in the introduction, Kamijo and Takeda (2010) found physical activity related differences in mixing costs dependent on the working memory demands of their two tasks. Despite our tasks being different to the tasks used by Kamijo and Takeda, it is interesting to see this same pattern of results in which there appears to be clear differences between exercise-related participant groups in conditions with the highest working memory demand.

These results are particularly interesting when we consider that some previous research has noted that high working memory demand is often associated with *decreased* vigilance (Caggiano & Parasuraman, 2004; Helton & Russell, 2011). However, this research has often been done with typical populations rather than populations with increased vigilance. We know from our previous research that martial artists have a higher level of vigilance than non-martial artists (Johnstone & Marí-Beffa, 2018), and therefore it may be that they still have an increased level of vigilance even with a small decrease due to the working memory demands. Therefore, it may be that the non-martial artists are more heavily affected by a decrease in vigilance caused by the working memory demands of the incongruent mixing costs. It is currently unclear if this is the case, and therefore future research considering the effects of Martial Arts experience on task switching may wish to include a measure of working memory capacity.

One way that we could get further clarification on whether these results are being driven by an increase in vigilance, would be to test participants on a random version of a task-switching paradigm (i.e., not a predictable pattern of trials) as well as an alternating runs version. As discussed in this paper’s introduction, the alternating runs paradigm involves an element of predictability for participants, which allows us to assess vigilance. If martial artists are performing at a greater level because of increased vigilance, we would only expect to see this greater performance in the alternating runs version of the task, and not the random version. Interestingly, Koch (2005) discussed how the predictability of the alternating runs paradigm is only beneficial if participants can hold the pattern in their working memory. They ran an experiment in which the predictability of the trials was manipulated, and they found that removing the predictability had a large impact on mixing costs due to the index’s reliance on repeat trials. Koch noted that due to the lack of predictability, participants were not receiving as much of a benefit from the repeat trials and were instead treating all the trials as potential switch trials.

The results from this paper support our lab’s previous finding that Martial Arts practice is associated with increased levels of vigilance. Given the differences in tasks (the Attention Network Test in Johnstone and Marí-Beffa (2018), and task-switching here), the fact that these previous findings have been replicated within different participants increases the robustness of the results. Indeed, despite not reaching the threshold for significance, the correlation between mixing costs (measuring vigilance) and years of Martial Arts experience, closely resembles that of the correlation that we previously found between the alert attentional network (also representing vigilance) and years of experience. While neither of these correlations reached significance, they both showed a trend towards significance, highlighting an interesting potential avenue for further study, especially with a larger *n* study featuring a wider range of years of experience.

With regards to switch costs, there was no significant difference between the martial artists and non-martial artists. This was surprising due to previous research suggesting a relationship between physical activity and cognitive flexibility. However, we should consider that the Kamijo and Takeda (2010) research on active vs sedentary populations stated that there may be more to uncover about a relationship between switch costs and physical activity, as their findings were only present in their behavioural results and not their electrophysiological data.

This paper aimed to use a task-switching paradigm to further investigate the effect of Martial Arts practice on vigilance, whilst also assessing cognitive flexibility. The results showed a smaller mixing cost in incongruent conditions for martial artists in comparison to non-martial artists, which we believe reflects an increased vigilance. We also found no significant difference in switch costs between martial artists and non-martial artists, suggesting that martial artists may not benefit from an increase in cognitive flexibility. We believe these results are important for informing research on which aspects of cognition may be able to be improved through Martial Arts practice.

## Acknowledgements

This research was funded by the Economic and Social Research Council Doctoral Training Centre (ESRC DTC) from United Kingdom, grant number ES/J500197/1.

## References

Caggiano, D. M., & Parasuraman, R. (2004). The role of memory representation in the vigilance decrement. Psychonomic bulletin & review, 11(5), 932–937.

Dajani, D. R., & Uddin, L. Q. (2015). Demystifying cognitive flexibility: Implications for clinical and developmental neuroscience. Trends in neurosciences, 38(9), 571–578.

Dreisbach, G., Haider, H., & Kluwe, R. H. (2002). Preparatory processes in the task-switching paradigm: Evidence from the use of probability cues. *Journal of Experimental Psychology: Learning*, Memory, and Cognition, 28(3), 468.

Helton, W. S., & Russell, P. N. (2011). Working memory load and the vigilance decrement. Experimental Brain Research, 212(3), 429–437.

Ionescu, T. (2012). Exploring the nature of cognitive flexibility. New ideas in psychology, 30(2), 190–200.

Johnstone, A., & Marí-Beffa, P. (2018). The Effects of Martial Arts Training on Attentional Networks in Typical Adults. Frontiers in Psychology, 9(80). doi:10.3389/fpsyg.2018.00080

Kamijo, K., & Takeda, Y. (2010). Regular physical activity improves executive function during task switching in young adults. International Journal of Psychophysiology, 75(3), 304–311.

Kirkham, A. J., Breeze, J. M., & Mari-Beffa, P. (2012). The impact of verbal instructions on goal-directed behaviour. Acta Psychol (Amst), 139(1), 212–219. doi:10.1016/j.actpsy.2011.09.016

Koch, I. (2005). Sequential task predictability in task switching. Psychonomic bulletin & review, 12(1), 107–112.

Lo, W. L. A., Liang, Z., Li, W., Luo, S., Zou, Z., Chen, S., & Yu, Q. (2019). The effect of judo training on set-shifting in school children. BioMed research international, 2019.

Marí-Beffa, P., & Kirkham, A. (2014). The mixing cost as a measure of cognitive control. Task Switching and Cognitive Control, 74–100.

Martin, M. M., & Rubin, R. B. (1995). A new measure of cognitive flexibility. Psychological reports, 76(2), 623–626.

Masley, S., Roetzheim, R., & Gualtieri, T. (2009). Aerobic exercise enhances cognitive flexibility. Journal of clinical psychology in medical settings, 16(2), 186–193.

Monsell, S. (2003). Task switching. Trends in Cognitive Sciences, 7(3), 134–140.

Rogers, R. D., & Monsell, S. (1995). Costs of a predictable switch between simple cognitive tasks. J Exp Psychol Gen, 124. doi:10.1037/0096-3445.124.2.207

Ruthruff, E., Remington, R. W., & Johnston, J. C. (2001). Switching between simple cognitive tasks: The interaction of top-down and bottom-up factors. Journal of Experimental Psychology: Human Perception and Performance, 27(6), 1404.

Sanabria, D., Luque-Casado, A., Perales, J. C., Ballester, R., Ciria, L. F., Huertas, F., & Perakakis, P. (2019). The relationship between vigilance capacity and physical exercise: a mixed-effects multistudy analysis. PeerJ, 7, e7118.

Sanchis, C., Blasco, E., Luna, F. G., & Lupiáñez, J. (2020). Effects of caffeine intake and exercise intensity on executive and arousal vigilance. Scientific reports, 10(1), 1–13.

